# Long-range inhibition synchronizes and updates prefrontal task activity

**DOI:** 10.1101/2022.03.31.486626

**Authors:** Kathleen K.A. Cho, Jingcheng Shi, Vikaas S. Sohal

**Author notes:** Institut Du Cerveau-Paris Brain Institute, Sorbonne Université, Inserm U1127, 47 Boulevard de L’Hôpital, 75013, Paris, France (Kathleen K.A. Cho).

## Abstract

Changes in patterns of activity within the medial prefrontal cortex enable rodents, non-human primates, and humans to update their behavior to adapt to changes in the environment, e.g., during cognitive tasks^1–5^. Within medial prefrontal cortex, inhibitory neurons expressing parvalbumin are important for updating strategies in a rule shift task^6–8^. Nevertheless, causal mechanisms through which parvalbumin neurons recruit specific circuits to produce prefrontal network dynamics that switch from maintaining to updating task-related patterns of activity remain unknown. Here we identify a new long-range inhibitory projection from prefrontal parvalbumin neurons to the contralateral cortex that mediates this process. Whereas nonspecifically inhibiting all callosal projections does not prevent mice from learning rule shifts, selectively inhibiting these callosal parvalbumin projections disrupts rule shift learning, desynchronizes gamma-frequency activity that is necessary for learning^8^, and suppresses changes in prefrontal activity patterns that normally accompany rule shifts. Thus, callosal parvalbumin projections switch prefrontal circuits from maintaining to updating behavioral strategies by synchronizing callosal communication and preventing it from inappropriately maintaining outdated neural representations. These findings may explain how deficits in prefrontal parvalbumin neurons and gamma synchrony cause impaired behavioral flexibility in schizophrenia^9–10^, and identify long-range projections from prefrontal parvalbumin neurons as a key circuit locus for understanding and treating these deficits.

## Main

Organisms must continually update their behavioral strategies to adapt to changes in the environment. Inappropriate perseveration on outdated strategies is a hallmark of conditions such as schizophrenia and bipolar disorder, and classically manifests in the Wisconsin Card Sorting Task (WCST)^11^. It is well documented that the prefrontal cortex is responsible for such flexible cognitive control, by providing active maintenance of rule or goal representations^12–14^, adaptive gating of these representations^15^, and the top-down biasing of sensory processing via extensive interconnectivity with other brain regions^16–17^. Studies have shown that within the medial prefrontal cortex (mPFC), normal parvalbumin (PV) interneuron function is required for mice to perform ‘rule shift’ tasks which, like the WCST, involve identifying uncued rule changes and learning new rules which utilize cues that were previously irrelevant to trial outcomes^6–7^. Moreover, PV interneurons play a key role in generating synchronized rhythmic activity in the gamma-frequency (∼40 Hz) range^18–19^. Indeed, during rule shift tasks, the synchrony of gamma-frequency activity between PV interneurons in the left and right mPFC increases after error trials, i.e., when mice receive feedback that a previously-learned rule has become outdated, and optogenetically disrupting this synchronization causes perseveration^8^. Nevertheless, the basic relationships between circuits (synaptic connections), network dynamics (interhemispheric gamma synchrony), and neural representations (task-dependent changes in activity patterns) remain unknown.

### Prefrontal parvalbumin neurons give rise to GABAergic callosal projections

Gamma synchrony is commonly assumed to be transmitted across regions by excitatory synapses^20^, which are the predominant form of long-range communication in the cortex. However, we explored an alternative hypothesis suggested by recent descriptions of long-range GABAergic connections originating from PV neurons in mPFC^21^. Specifically, we first visualized that PV-expressing neurons in the mPFC give rise to callosal GABAergic synapses in the contralateral mPFC (Fig. 1). We identified this anatomical link by injecting AAV-EF1α-DIO-ChR2-eYFP into one mPFC of *PV-Cre* mice and observed virally-labeled PV terminals in the contralateral PFC, particularly in deep layers 5 and 6 (Fig. 1a). To characterize this novel callosal PV projection and its recipient neurons, we performed recordings in the contralateral mPFC (Fig. 1b). It has been reported that mPFC pyramidal cells are not homogeneous, but rather can be grouped by their projection targets with corresponding anatomical and electrophysiological properties^22^ (Fig. 1c). Our recordings reveal that callosal PV projections preferentially inhibit the ‘pyramidal tract’ (PT) pyramidal neuron subtype (21/31 connected), which projects subcortically to thalamus or brainstem, compared to the ‘intratelencephalic’ (IT) subtype (10/44 connected), which projects to contralateral cortex and/or striatum (Fig. 1d). Furthermore, rhythmic trains of PV-terminal optogenetic stimulation elicited time-locked inhibitory postsynaptic potentials (IPSPs) in ChR2-negative pyramidal neurons (Fig. 1e). These optogenetically-evoked inhibitory postsynaptic potentials (IPSPs) were not affected by blocking excitatory synapses using glutamatergic receptor antagonists CNQX and APV (10 μM and 50 μM, respectively), but were completely abolished by bath application of the GABA_A_ receptor antagonist gabazine (10 μM; Fig.1f).

**Fig. 1.**
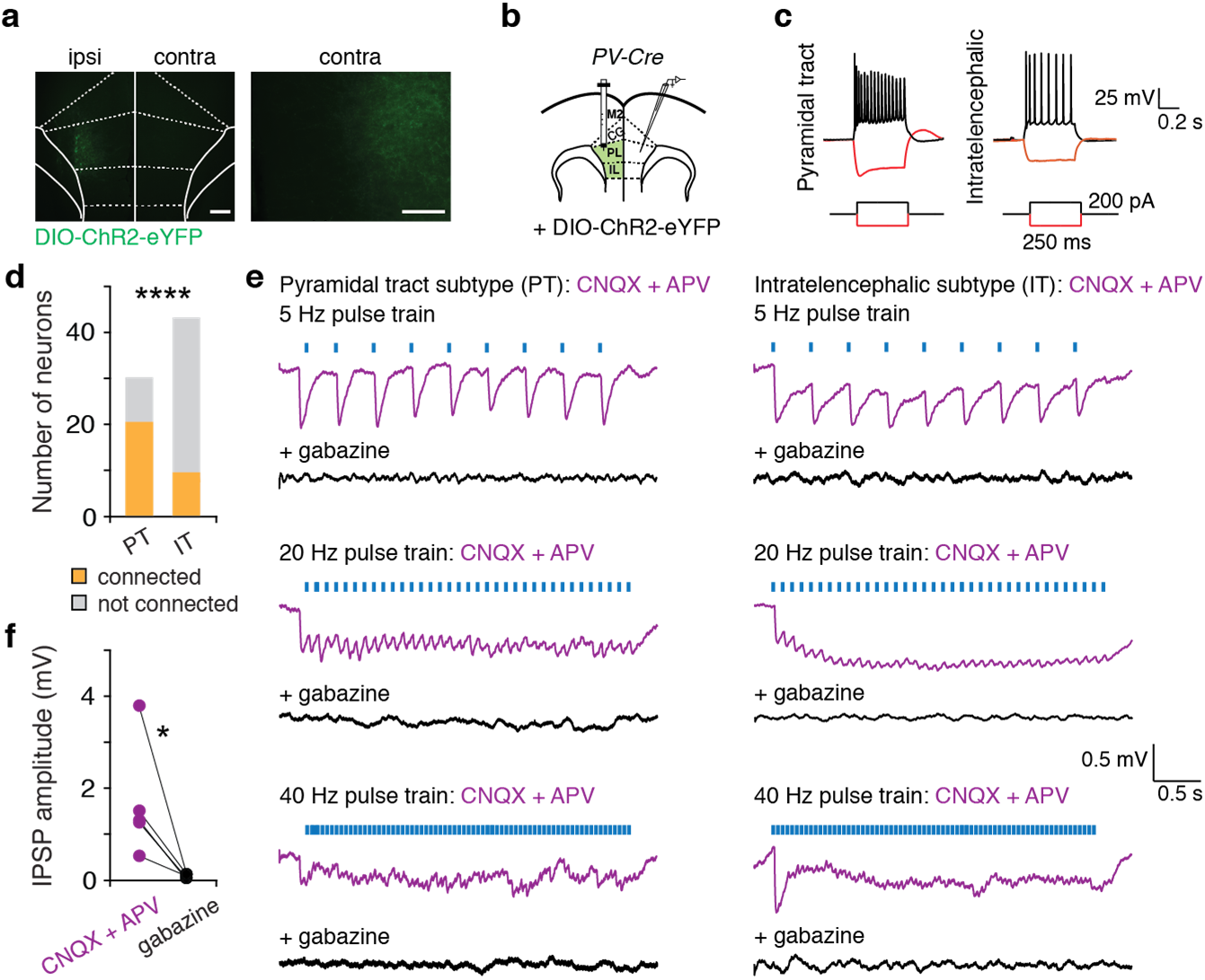
Callosal mPFC PV projections preferentially target pyramidal tract neurons. **a**, Injection of AAV-DIO-ChR2-eYFP in one (ipsi) mPFC of *PV-Cre* mice enabled visualization of eYFP+ terminals in the contralateral (contra) mPFC. **b**, Experimental design schematic. Whole-cell recordings were made contralateral to the virus injection in prefrontal brain slices. **c**, Example current-clamp responses from recipient neurons during injection of hyperpolarizing or depolarizing current. **d**, During optogenetic stimulation of callosal PV terminals, we observed synaptic responses in a greater fraction of pyramidal tract (PT) neurons compared to the intratelencephalic (IT) subtype (Chi-square test, *P* < 0.0001; *n* = 75 cells, 11 mice). **e**, Rhythmic trains of blue light flashes (5 ms, 470 nm; denoted as blue bars) delivered through the 40x objective were used to optogenetically stimulate callosal PV terminals. Example recordings from PT and IT subtypes showing optogenetically-evoked inhibitory postsynaptic potentials (IPSPs) in the presence of CNQX + APV that are abolished in gabazine. **f**, IPSPs are blocked by gabazine application (two-tailed paired *t-*test, *t*_(4)_ = 2.953, *P* = 0.0419; *n* = 5 cells, 4 mice). \**P* < 0.05, \*\*\*\**P* < 0.0001; scale bars, 250 μm and 100 μm, respectively.

### Callosal PV^+^ connections are necessary for rule shifts

While we identify a long-range inhibitory anatomical connection between the prefrontal cortices, the function of this input is not defined. We next explored the role of these callosal mPFC PV^+^ projections as mice perform a task involving the type of behavioral adaptations involved in the WCST. Variants of this task have previously been characterized by our laboratory and others^6–7,23^ in which mice are first required to associate one set of sensory cues with the location of a hidden food reward (initial association). After learning the initial association, mice must then learn to attend to a set of sensory cues that were previously present, but irrelevant to the outcome of each trial, and make decisions based on these previously-irrelevant cues (rule shift) (Fig. 2a, Extended Data Fig. 1). Because cross-hemispheric gamma synchrony between PV interneurons is essential for learning the rule shift^8^, callosal PV^+^ projections may be a good candidate to effectively coordinate activity across hemispheres. To investigate this, we injected AAV-EF1α-DIO-eNpHR-mCherry or control virus in one mPFC of *PV-Cre* mice, then optogenetically-silenced callosal PV^+^ terminals in the contralateral mPFC during the rule shift portion of the task (Fig. 2b–f, Extended Data Fig. 1b, and Extended Data Fig. 2). Selective inhibition of callosal PV^+^ terminals is sufficient to impair rule shift learning and induce perseveration (relative to *PV-Cre* mice injected with a control virus; Fig. 2e–f, Extended Data Fig. 2a–h). Interestingly, non-specific inhibition of all callosal projections (both GABAergic and excitatory connections), did not affect rule-shift performance (Fig. 2g–j, Extended Data Fig. 2i–p). These results demonstrate that callosal PV^+^ projections are instrumental for proper communication across hemispheres, such that suppressing them causes aberrant interhemispheric communication which disrupts normal learning. In particular, long-range GABAergic projections have been hypothesized to play a role in the temporal coordination of neuronal activity^24–25^, suggesting that when callosal interactions are restricted to excitatory connections, communication between the two hemispheres may become mistimed.

**Fig. 2.**
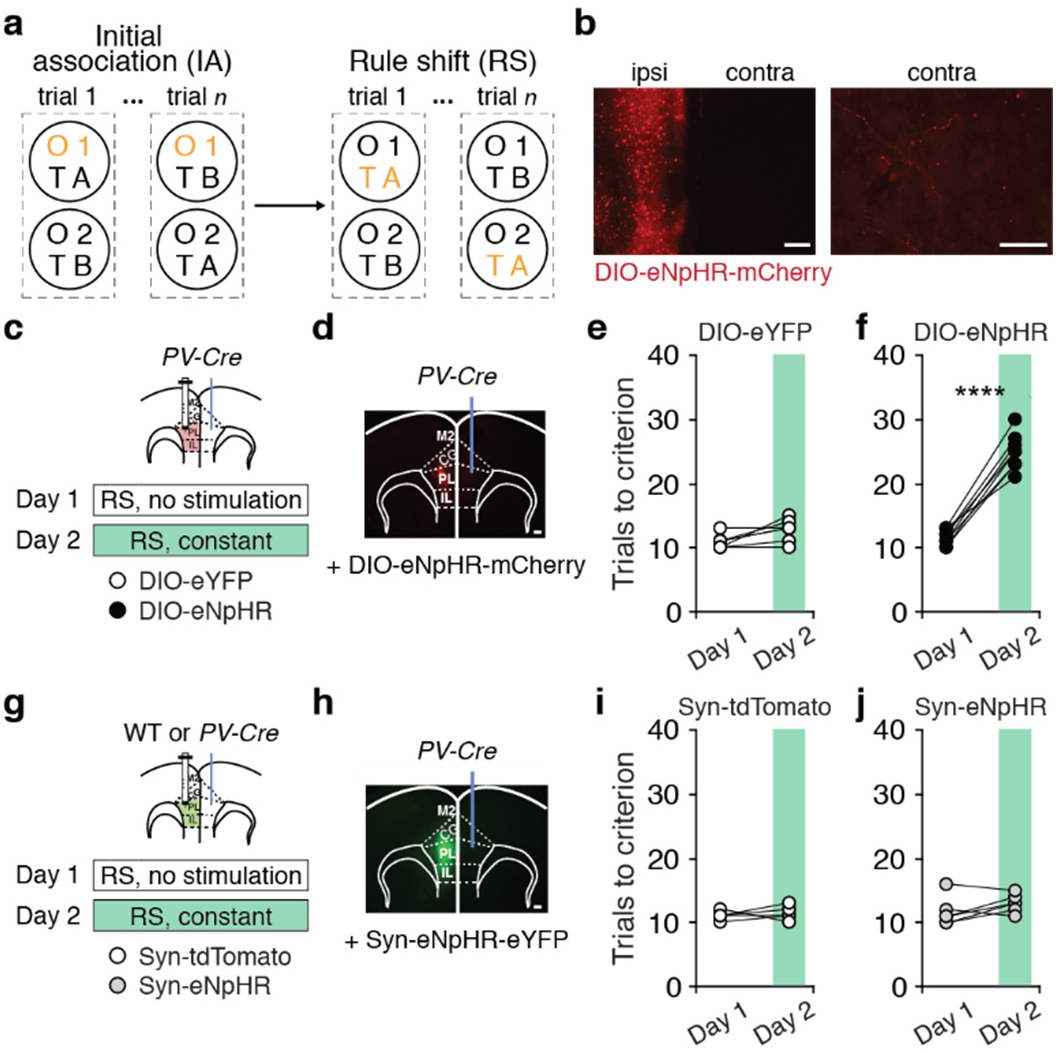
Optogenetic inhibition of callosal mPFC PV projections impairs cognitive flexibility. **a**, Rule shift task schematic. On each trial, a mouse chooses one of two bowls, each scented with a different odor (O1 or O2) and filled with a different textured digging medium (TA or TB), to find a food reward. Mice first learn an initial association (IA) between one of these sensory cues (for example, odor O1) and food reward (the cue associated with reward is indicated in orange). Once mice reach the learning criterion (8 of 10 consecutive trials correct), this association undergoes an extra-dimensional rule shift (RS; for example, from O1 to TA being rewarded). **b**, Representative image showing AAV-DIO-eNpHR-mCherry (DIO-eNpHR) expression in ipsilateral mPFC (ipsi) and callosal PV terminals in contralateral mPFC (contra; scale bars, 100 μm and 50 μm, respectively). **c, g**, Experimental design: Day 1, no light delivery; Day 2, continuous light for inhibition during the rule shift (RS). **d**, Representative image showing DIO-eNpHR expression in one mPFC and a fiber-optic cannula implanted in the contralateral mPFC. Scale bar, 100 μm. **e, f**, Optogenetic inhibition of mPFC callosal PV terminals impairs rule shift performance in DIO-eNpHR-expressing mice (*n* = 8) compared to controls (*n* = 7) (two-way ANOVA (task day × virus); interaction: *F*_(1,13)_ = 16.1, *P* = 0.0015). **e**, Performance of DIO-eYFP-expressing controls did not change from Day 1 to Day 2 (post hoc *t*_(13)_ = 1.273, *P* = 0.4508). **f**, Inhibition disrupts rule shift performance in DIO-eNpHR-expressing mice (post hoc *t*_(13)_ = 7.235, *P* < 0.0001). **h**, Representative image showing AAV-Synapsin-eNpHR-eYFP (Syn-eNpHR) expression in one mPFC and a fiber-optic cannula implanted in the contralateral mPFC. Scale bar, 100 μm. **i, j**, Performance of Synapsin-tdTomato-expressing controls (Syn-tdTomato, *n* = 4) is similar to Syn-eNpHR-expressing mice (*n* = 6) from Day 1 to Day 2 (two-way ANOVA (task day × virus); interaction: *F*_(1,8)_ = 0.01553, *P* = 0.9039). Two-way ANOVA followed by Bonferroni post hoc comparisons was used. \*\*\*\**P* < 0.0001.

### Callosal PV^+^ connections are necessary for increases in gamma synchrony during rule shifts

To test this possibility, we studied how callosal PV^+^ projections influence the cross-hemispheric gamma synchrony in PV interneurons that we previously found is essential for re-appraising the behavioral salience of sensory cues during rule shifts. For this, we injected AAV-EF1α-DIO-eNpHR-BFP or control virus in one mPFC of *PV-Cre* Ai14 mice (Fig. 3a, Extended Data Fig. 3a–b). To track transmembrane voltage activity patterns of PV neurons, we also injected the Cre-dependent, genetically-encoded voltage indicator AAV-DIO-Ace2N-4AA-mNeon (Ace-mNeon) in both mPFCs. We then implanted multimode optical fibers to excite and measure fluorescence from both Ace-mNeon and a control fluorophore (tdTomato) expressed in PV interneurons in the left and right mPFC as well as optogenetically-silence PV callosal terminals (Fig. 3a–b, Extended Data Fig. 1c). Using this method, we had previously found that gamma synchrony between PV interneurons in the left and right mPFC increases during rule shifts when mice make errors, i.e., when they fail to receive an expected reward and thereby receive feedback that the previously learned association is no longer valid^8^. We confirmed this result on Day 1 (Fig. 3c–g), in the absence of any optogenetic silencing. On Day 2, optogenetic inhibition of callosal PV^+^ terminals disrupted performance in the rule shift task, compared to *PV-Cre* Ai14 mice given control virus (Fig. 3c–e, Extended Data Fig. 3). This behavioral deficit, which was consistent with our earlier experiments, correlated with deficits in cross-hemispheric gamma synchrony: specifically, the increase in cross-hemispheric PV neuron gamma synchrony observed after rule shift errors on Day 1, was abolished on Day 2 (Fig. 3f–g). Surprisingly, the impairments in both rule shift performance and gamma synchrony persisted on Day 3, in the absence of any additional optogenetic inhibition (Fig. 3e, g). These data show that callosal PV^+^ projections normally promote the interhemispheric gamma synchrony that we previously found was necessary for rule shifts. Thus, when callosal PV^+^ projections are selectively suppressed, communication between the hemispheres becomes mistimed, potentially affecting the normal evolution of prefrontal network activity during rule shifts.

**Fig. 3.**
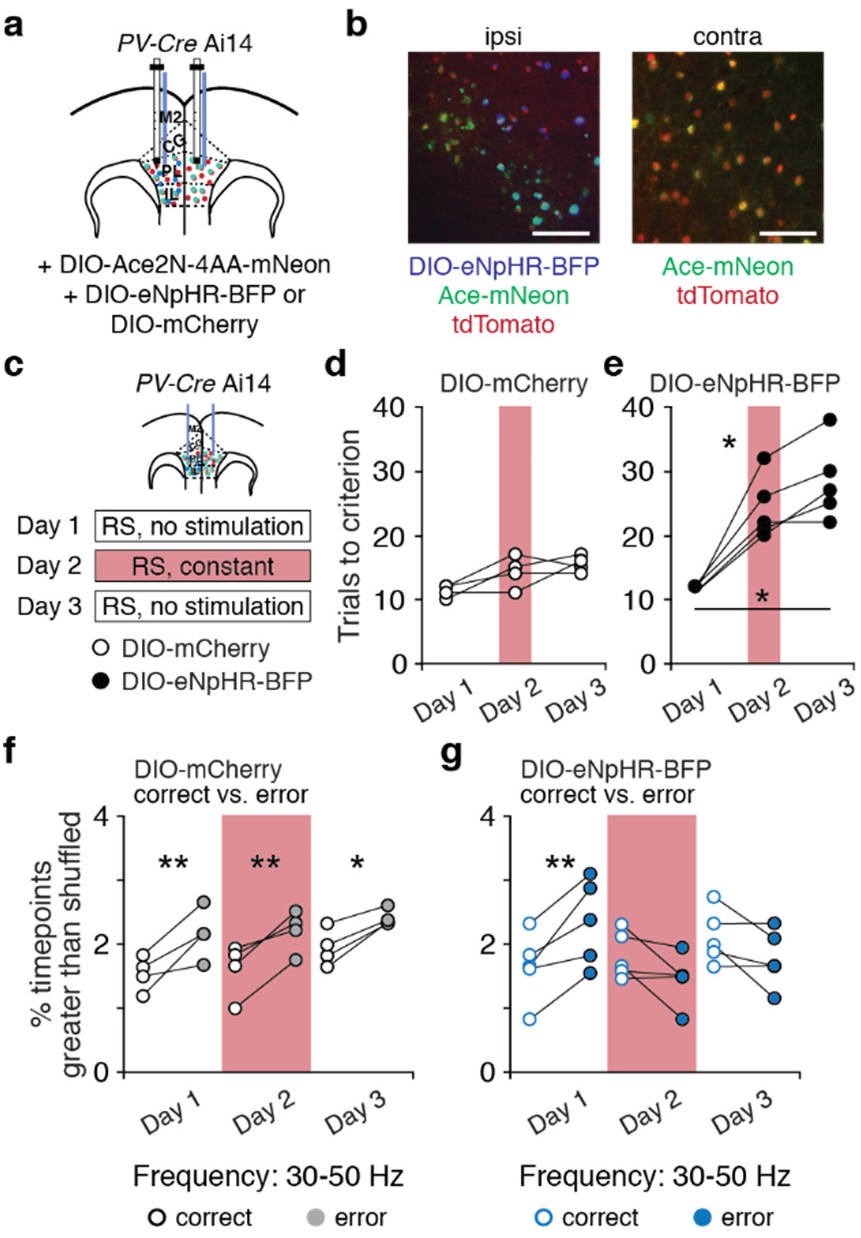
Optogenetic inhibition of callosal mPFC PV projections disrupts interhemispheric gamma synchrony. **a**, *PV-Cre* Ai14 mice had bilateral AAV-DIO-Ace2N-4AA-mNeon (Ace-mNeon) injections, an ipsilateral AAV-DIO-eNpHR-BFP (DIO-eNpHR) or control virus (AAV-DIO-mCherry) injection and multimode fiber-optic implants in both prefrontal cortices. **b**, Representative image showing DIO-eNpHR, Ace-mNeon, and tdTomato expression in the ipsilateral mPFC (ipsi) and Ace-mNeon and tdTomato in the contralateral hemisphere (contra). **c**, Experimental design: Day 1, no light delivery; Day 2, continuous light for inhibition during the rule shift (RS); Day 3, no light delivery. **d, e**, Optogenetic inhibition of callosal mPFC PV terminals impairs rule shift performance in DIO-eNpHR-expressing mice (*n* = 5) compared to DIO-mCherry-expressing controls (*n* = 4) on Day 2 and Day 3 (two-way ANOVA (task day × virus); interaction: *F*_(2,14)_ = 11.98, *P* = 0.0009). **d**, Performance of DIO-mCherry-expressing controls did not change from Day 1 to Day 3 (Day 1 to Day 2: post hoc *t*_(3)_ = 2.449, *P* = 0.2752; Day 1 to Day 3: post hoc *t*_(3)_ = 3.833, *P* = 0.0939; Day 2 to Day 3: post hoc *t*_(3)_ = 0.8372, *P* > 0.9999). **e**, Optogenetic inhibition of mPFC callosal PV terminals impairs rule shift performance in DIO-eNpHR-expressing mice compared to DIO-mCherry-expressing controls (Day 1 to Day 2: post hoc *t*_(4)_ = 5.759, *P* = 0.0135; Day 1 to Day 3: post hoc *t*_(4)_ = 6.010, *P* = 0.0116; Day 2 to Day 3: post hoc *t*_(4)_ = 3.5, *P* = 0.0747). **f**, Cross-hemispheric 30-50 Hz synchronization is higher after RS errors than after RS correct decisions across task days in control mice (two-way ANOVA; Day 1: post hoc *t*_(9)_ = 4.695, *P* = 0.0034; Day 2: post hoc *t*_(9)_ = 4.415, *P* = 0.005; Day 3: post hoc *t*_(9)_ = 3.455, *P* = 0.0216). **g**, While cross-hemispheric gamma synchrony is higher after RS errors than after RS correct decisions for DIO-eNpHR-expressing mice with no light delivery on Day 1, it was not different after errors vs. after correct decisions on Days 2 and 3 (two-way ANOVA; Day 1: post hoc *t*_(12)_ = 3.97, *P* = 0.0056; Day 2: post hoc *t*_(12)_ = 2.127, *P* = 0.1646; Day 3: post hoc *t*_(12)_ = 1.936, *P* = 02302). Two-way ANOVA followed by Bonferroni post hoc comparisons was used. *\*P* < 0.05, \*\**P* < 0.01; scale bars, 50 μm.

### Inhibiting callosal PV^+^ connections disrupts the reorganization of prefrontal activity during rule shifts

We specifically hypothesized that the inhibition of callosal PV^+^ projections and consequent deficits in gamma synchrony would disrupt the changes in prefrontal activity patterns that normally serve to update behavioral strategies^3^. In particular, gamma synchrony normally increases specifically following rule shift errors. Therefore, we hypothesized that prefrontal activity patterns might begin to diverge from previously-established representations specifically during this period, and that inhibiting callosal PV^+^ projections would disrupt this process. To quantify this, we measured the similarity between activity patterns occurring during the 10 seconds immediately following a choice on correct versus error trials. We measured these activity patterns using microendoscopic calcium imaging in *PV-Cre* mice injected with AAV-Synapsin-GCaMP7f (Syn-GCaMP7f) and implanted with a gradient-index (GRIN) lens in one mPFC, and injected with AAV-EF1α-DIO-eNpHR-mCherry or a control virus in the other mPFC (Fig. 4a–b, Extended Data Fig. 4a–b). We measured Syn-GCaMP7f fluorescence while mice performed a rule shift on three consecutive days (Fig. 4a). On the second day, we also delivered red light to activate eNpHR in callosal PV^+^ terminals (Extended Data Fig. 1d). As in previous experiments, inhibiting callosal PV^+^ terminals significantly increased the number of trials needed to learn a rule shift in eNpHR-expressing mice (whereas there was no effect in controls) (Fig. 4c–d, Extended Data Fig. 4). This was accompanied by a significant increase in the similarity between activity patterns occurring after error trials and those after correct trials (by contrast, no such increase occurred in controls) (Fig. 4e–f). To confirm that this reflects a suppression of changes in activity patterns that normally occur after rule shift errors, we also computed the similarity between activity patterns after rule shift errors and those occurring after errors during learning of the initial association (Fig. 4g–f). Activity patterns following rule shift errors became more similar to those observed during the initial association. Thus, inhibiting callosal PV^+^ projections prevents prefrontal activity patterns from diverging from previously-established representations.

**Fig. 4.**
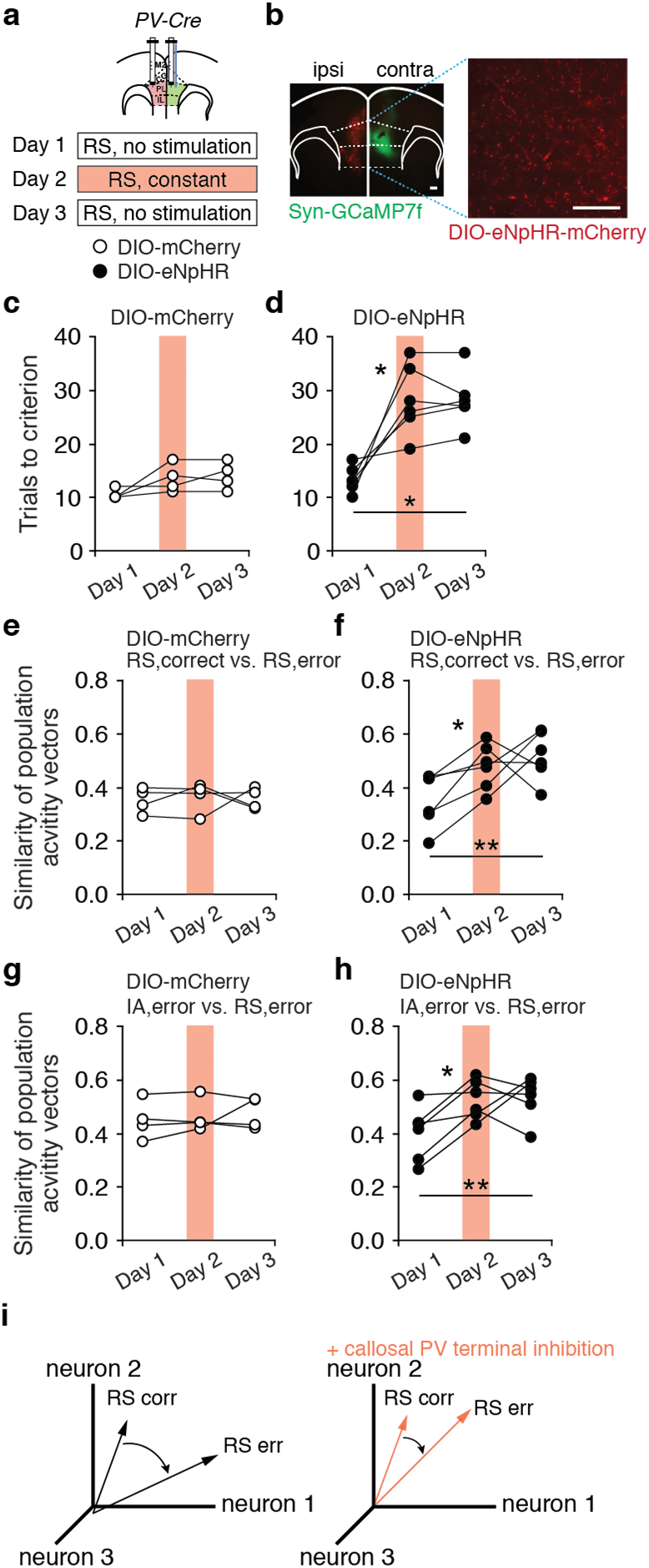
Optogenetic inhibition of callosal mPFC PV projections changes prefrontal activity patterns. **a**, *PV-Cre* mice had AAV-DIO-eNpHR-mCherry (DIO-eNpHR) or control AAV-DIO-mCherry (DIO-mCherry) injected in the ipsilateral hemisphere (ipsi), AAV-Synapsin-GCaMP7f (Syn-GCaMP7f) injected in the contralateral hemisphere (contra), and a GRIN lens connected to a miniscope implanted in the contra hemisphere. Experimental design: Day 1, no light delivery; Day 2, continuous light for inhibition during the rule shift (RS); Day 3, no light delivery. **b**, Left: Representative image showing DIO-eNpHR injected in the ipsi mPFC and Syn-GCaMP7f in the contra hemisphere. Right: DIO-eNpHR expression in callosal PV axonal fibers. **c, d**, Optogenetic inhibition of callosal mPFC PV terminals impairs rule shift performance in DIO-eNpHR mice (*n* = 6) compared to DIO-mCherry-expressing controls (*n* = 4) on Day 2 and Day 3 (two-way ANOVA (task day × virus); interaction: *F*_(2,16)_ = 6.882, *P* = 0.007). **c**, Performance of DIO-mCherry-expressing controls did not change from Day 1 to Day 3 (Day 1 to Day 2: post hoc *t*_(3)_ = 1.897, *P* = 0.4621; Day 1 to Day 3: post hoc *t*_(3)_ = 2.782, *P* = 0.2067; Day 2 to Day 3: post hoc *t*_(3)_ = 0.5774, *P* > 0.9999). **d**, Optogenetic inhibition of mPFC callosal PV terminals impairs rule shift performance in DIO-eNpHR-expressing mice (Day 1 to Day 2: post hoc *t*_(5)_ = 4.174, *P* = 0.0261; Day 1 to Day 3: post hoc *t*_(5)_ = 4.901, *P* = 0.0134; Day 2 to Day 3: post hoc *t*_(5)_ = 0.000, *P* > 0.9999). **e, f**, Optogenetic inhibition changes the similarity of population activity vectors (two-way ANOVA, main effect of virus: *F*_(1,8)_ = 8.788, *P* = 0.018). **e**, For control mice, there is no change in the similarity between population activity vectors after RS errors and those after correct decisions across task days (Day 1 to Day 2: post hoc *t*_(16)_ = 0.2349, *P* > 0.9999; Day 1 to Day 3: post hoc *t*_(16)_ = 0.1156, *P* > 0.9999; Day 2 to Day 3: post hoc *t*_(16)_ = 0.1193, *P* > 0.9999). **f**, In DIO-eNpHR-expressing mice, there is an increase in the similarity between population activity vectors after RS errors and those after correct decisions from Day 1 (before optogenetic inhibition) to Days 2 and 3 (during and after inhibition, respectively) (Day 1 to Day 2: post hoc *t*_(16)_ = 2.802, *P* = 0.0384; Day 1 to Day 3: post hoc *t*_(16)_ = 3.675, *P* = 0.0061; Day 2 to Day 3: post hoc *t*_(16)_ = 0.8738, *P* > 0.9999). **g, h**, We computed the similarity between population activity vectors observed after decisions on error trials during the IA and those during the RS. There is increased similarity across days in DIO-eNpHR-expressing mice (*n* = 6) (two-way ANOVA; main effect of task day: *F*_(2,16)_ = 4.538, *P* = 0.0275). **g**, There is no change in the similarity of population activity vectors after initial association (IA) errors and RS errors across task Days in control mice (*n* = 4; Day 1 to Day 2: post hoc *t*_(16)_ = 0.3205, *P* > 0.9999; Day 1 to Day 3: post hoc *t*_(16)_ = 0.609, *P* > 0.9999; Day 2 to Day 3: post hoc *t*_(16)_ = 0.2885, *P* > 0.9999). **h**, During and following optogenetic inhibition on Days 2 and 3 in eNpHR-expressing mice, there is an increase in the similarity of population activity vectors after between IA and RS error trials (Day 1 to Day 2: post hoc *t*_(16)_ = 3.424, *P* = 0.0104; Day 1 to Day 3: post hoc *t*_(16)_ = 3.631, *P* = 0.0067; Day 2 to Day 3: post hoc *t*_(16)_ = 0.2066, *P* > 0.9999). **i**, Schematic illustrating the increase in the similarity of population activity vectors between RS correct and error trials with and without callosal PV terminal inhibition. Two-way ANOVA followed by Bonferroni post hoc comparisons was used. *\*P* < 0.05, \*\**P* < 0.01; scale bars, 100 μm and 50 μm, respectively.

As in previous experiments, deficits in learning a rule shift observed after inhibiting callosal PV^+^ projections persisted on the next day (‘Day 3’) without any additional light delivery to activate eNpHR (Fig. 4d). The increased similarity between activity patterns following rule shift errors and those following correct trials and between rule shift errors and initial association errors also persisted at this later time point (Fig. 4f, h). By contrast, there was no change in the similarity of activity patterns after correct versus error rule shift trials across days in control (eNpHR-negative) mice (Fig. 4e).

## Discussion

Changes in patterns of activity within the prefrontal cortex are presumed to drive behavioral adaptation. But the neural mechanisms which switch the mPFC from maintaining previously learned representations and strategies to updating them have not been known. Previous work had suggested that long-distance gamma synchrony might play a role. This study not only shows that impairments in gamma synchrony are indeed associated with impaired evolution of prefrontal activity patterns and the inappropriate maintenance of previously-learned representations, but reveals three surprising features. First, callosal GABAergic synapses originating from prefrontal PV neurons are required for learning rule shifts and generate increases in gamma synchrony that occur during learning. The fact that neocortical GABAergic neurons give rise to long-range connections capable of eliciting physiologically meaningful postsynaptic responses has only recently become appreciated, and the behavioral functions of these long-range GABAergic projections are not well-understood. Here we show that in mPFC, callosal GABAergic projections from PV^+^ neurons promote *inter*hemispheric gamma synchrony which has previously been shown to be necessary for learning rule shifts. This directly supports a hypothesized role of long-range GABAergic projections in synchronizing rhythmic activity interhemispherically^24^, and contrasts with earlier findings from hippocampal slices which suggested that excitatory synapses on PV interneurons may synchronize gamma oscillations across long distances^26^. In particular, whereas many long-range GABAergic projections mainly target downstream GABAergic neurons^22^, callosal PV^+^ projections innervate a large fraction of layer V pyramidal neurons. Notably, callosal PV^+^ synapses have the same subtype-specificity previously observed for local PV interneuron synapses^27^, in that they preferentially inhibit the PT, rather than the IT, subtype of layer 5 pyramidal cells.

Second, in this task, gamma synchrony does not appear to act via the ‘communication through coherence’ (CTC) mechanism^27^. CTC posits that when two regions are synchronized in a coherent manner, they are able to communicate constructively. However, in this case, communication between the two hemispheres is not actually necessary for rule shift learning, as mice learn normally even when we nonspecifically inhibit all callosal projections. By contrast, when PV^+^ callosal projections are selectively inhibited, sparing excitatory callosal communication, mice become perseverative and previously-learned representations are inappropriately maintained. In this scenario, excitatory communication between the two hemispheres is intact, but gamma synchrony is lost (Fig. 3g). We previously showed that optogenetically perturbing gamma synchrony (using out-of-phase stimulation across hemispheres) disrupts rule shift learning^8^. Thus, inter-regional communication is not necessary for normal behavior; however, when gamma synchrony is lost, inter-regional communication occurs in an aberrant manner that interferes with normal learning. This could occur because, during a gamma cycle, high levels of inhibition gradually fall, producing corresponding changes in excitatory neuron firing^8^. In this way, rhythmic inhibition periodically resets network activity; this reset could allow the emergence of new representations which diverge from the previously-established ones that are stored within patterns of recurrent connections. By contrast, when excitatory input from the contralateral cortex is not appropriately synchronized by PV-driven inhibition, it could arrive prematurely, inappropriately reinforcing previously-learned representations and thereby disrupting normal learning.

A third surprising feature is that inhibiting callosal PV^+^ projections leads to persistent changes in gamma synchrony, neural representations, and rule shift performance. This represents a novel form of network plasticity, whereby transiently disrupting connectivity between two brain regions leads to persistent deficits in their ability to synchronize, process information, and contribute to behavior. Notably, the resulting network state, characterized by cognitive inflexibility and deficient task-evoked gamma synchrony, captures key cognitive and circuit endophenotypes of schizophrenia. In many respects, this deleterious network plasticity represents the inverse of our earlier finding that transiently *enhancing* gamma synchrony (using either synchronizing optogenetic stimulation or low doses of the benzodiazepine clonazepam) can persistently *rescue* rule shift learning in mutant (*Dlx5/6*^+/-^) mice ^6,8^. Our new findings show that prefrontal circuit plasticity resulting from transient modulations of gamma synchrony is bi-directional, occurs even in previously-normal mice, and can be triggered by a previously unappreciated long-range GABAergic projection.

In summary, our results reveal circuit mechanisms which switch the prefrontal cortex from maintaining to updating behavioral strategies, functions for interhemispheric GABAergic projections in the neocortex, roles for gamma synchrony in information processing beyond communication through coherence, and new forms of network plasticity. These findings identify a new callosal connection originating from prefrontal parvalbumin neurons as a critical circuit locus for understanding and modulating the deficits in gamma synchrony and behavioral flexibility that are major features of schizophrenia^9–10^.

## Methods

### Mice

All animal care, procedures, and experiments were conducted in accordance with the NIH guidelines and approved by the Administrative Panels on Laboratory Animal Care at the University of California, San Francisco. Mice were group housed (2-5 siblings) in a temperature-controlled environment (22-24°C), had *ad libitum* access to food and water, and reared in normal lighting conditions (12-h light-dark cycle), until rule shift experiments began. All experiments were done using *PV-Cre*, wild-type C57/Bl6, and *PV-Cre* Ai14 lines (The Jackson Laboratory). Both male and female adult mice (10-20 weeks at time of experiment) were used in the behavioral experiments.

### Surgery

Male and female mice were anaesthetized using isoflurane (2.5% induction, 1.2-1.5% maintenance, in 95% oxygen) and placed in a stereotaxic frame (David Kopf Instruments). Body temperature was maintained using a heating pad. An incision was made to expose the skull for stereotaxic alignment using bregma and lambda as vertical references. The scalp and periosteum were removed from the dorsal surface of the skull and scored with a scalpel to improve implant adhesion. Viruses were infused at 100-150 nL/min through a 35-gauge, beveled injection needle (World Precision Instruments) using a microsyringe pump (World Precision Instruments, UMP3 UltraMicroPump). After infusion, the needle was kept at the injection site for 5-10 min and then slowly withdrawn. After surgery, mice were allowed to recover until ambulatory on a heated pad, then returned to their home cage.

For slice electrophysiology experiments using optogenetic opsins, mice were injected unilaterally in the mPFC, near the border between the prelimbic and infralimbic cortices (1.7 anterior-posterior (AP), + or - 0.3 mediolateral (ML), and −2.75 dorsoventral (DV) millimeters relative to bregma) with 0.4 μL of AAV5-EF1α-DIO-ChR2-eYFP (UNC Virus Core) to selectively target neurons expressing Cre. To allow for virus expression, experiments began at least 3 weeks after injection.

For behavioral experiments using optogenetics, mice were injected unilaterally in the mPFC, near the border between the prelimbic and infralimbic cortices (1.7 anterior-posterior (AP), + or −0.3 mediolateral (ML), and −2.75 dorsoventral (DV) millimeters relative to bregma) with 1 μL of AAV2-EF1α-DIO-eNpHR3.0-mCherry (UNC Virus Core), 1 μL of AAV5-EF1α-DIO-eYFP (UNC Virus Core), AAV5-hSynapsin-eNpHR3.0-eYFP (UNC Virus Core) or AAV5-Synapsin-tdTomato (UNC Virus Core), to selectively target Cre-expressing cells or non-selectively target prefrontal neurons. After injecting virus, a 200/240 μm (core/outer) diameter, NA=0.22, mono fiber-optic cannula (Doric Lenses, DFC_200/240-0.22_2.3mm_FLT) was slowly inserted into mPFC until the tip of the fiber reached a DV depth of −2.25. Implants were affixed onto the skull using Metabond Quick Adhesive Cement (Parkell). To allow for virus expression, behavioral experiments began at least 5 weeks after injection.

For behavioral experiments that combined dual-site voltage indicator imaging with optogenetics, mice were injected bilaterally at 3 depths (DV: −2.5, −2.25, −2.0) at the following AP/ML for mPFC: 1.7 AP, ±0.3 ML with 3 × 0.2 μL of AAV1-CAG-DIO-Ace2N-4AA-mNeon (Virovek). Mice were also injected unilaterally in the mPFC (1.7 anterior-posterior (AP), −0.3 mediolateral (ML), and −2.75 dorsoventral (DV) millimeters relative to bregma) with either 1 uL of AAV2-EF1α-DIO-eNpHR3.0-BFP (Virovek) or 1 μL of AAV5-EF1α-DIO-mCherry (UNC Virus Core). After injection of virus, two 400/430 μm (core/outer) diameter, NA=0.48, multimode fiber implants (Doric Lenses, MFC_400/430-0.48_2.8mm_ZF1.25_FLT) were slowly inserted into the mPFC at a ±12° angle using the following coordinates: 1.7 (AP), ±0.76 (ML), −2.14 (DV). To allow for virus expression, behavioral experiments began at least 5 weeks after injection.

For behavioral experiments that combined *in vivo* calcium imaging with optogenetics, mice were injected bilaterally at 4 depths (DV: −2.75, −2.5, −2.25, −2.0) at the following AP/ML for mPFC: 1.7 AP, +0.4 ML with diluted (1:3; Addgene) 4 × 0.15 μL of AAV9-Synapsin-jGCaMP7f-WPRE (Addgene). Mice were also injected unilaterally in the mPFC (1.7 anterior-posterior (AP), −0.3 mediolateral (ML), and −2.75 dorsoventral (DV) millimeters relative to bregma) with either 1 uL of AAV2-EF1α-DIO-eNpHR3.0-mCherry (Virovek) or 1 μL of AAV5-EF1α-DIO-mCherry (UNC Virus Core). After injection of virus, a 0.5 mm x 4.0 mm long integrated GRIN lens (Inscopix) was slowly advanced into the mPFC until the tip was placed at 1.7 AP, +0.4 ML, DV −2.25 and cemented in place with Metabond Quick Adhesive Cement (Parkell). To allow for virus expression, behavioral experiments began at least 5 weeks after injection.

### Slice preparation and analysis

Adult mice were anesthetized with an intraperitoneal injection of euthasol, decapitated and the brains were rapidly removed. 250 μm thick coronal slices were cut from adult mice of either sex using a vibratome (VT1200S Leica Microsystems, Inc.) and a chilled slicing solution in which Na^+^ was replaced by sucrose, then incubated in warmed ACSF at 30-31°C for 15 minutes and then at least one hour at room temperature before being used for recordings. ACSF contained (in mM): 126 NaCl, 26 NaHCO3, 2.5 KCl, 1.25 NaH2PO4, 1 MgCl2, 2 CaCl, and 10 glucose. Slices were secured by placing a harp along the midline between the two hemispheres.

Somatic whole-cell patch recordings were obtained from ChR2-negative neurons amid ChR2-positive axonal fibers in the mPFC contralateral to the site of virus injection, on an upright microscope (BX51WI; Olympus). Recordings were made using a Multiclamp 700A (Molecular Devices). Patch electrodes (tip resistance = 2–6 MOhms) were filled with the following (in mM): 130 K-gluconate, 10 KCl, 10 HEPES, 10 EGTA, 2 MgCl, 2 MgATP, and 0.3 NaGTP (pH adjusted to 7.3 with KOH). All recordings were at 32.5±1°C. Series resistance was usually 10–20 MΩ, and experiments were discontinued above 30 MΩ.

Intrinsic properties were calculated based on the current clamp responses to a series of 250 msec current pulse injections from −200 to 450 pA (50 pA/increment). Spiking properties were calculated based on the response to a current pulse that was 100 pA above the minimal level that elicited spiking. Pyramidal tract neuron subtypes were identified based on characteristic firing patterns, specifically h-current-induced “sag” > 3mV in response to hyperpolarizing current pulses.

### *In vitro* ChR2 stimulation

Stimulation of channelrhodopsin (ChR2) in callosal PV terminals was performed using ∼4-5mW flashes of light generated by a Lambda DG-4 high-speed optical switch with a 300W Xenon lamp (Sutter Instruments), and an excitation filter set centered around 470 nm, delivered to the slice through a 40x objective (Olympus). Illumination was delivered across a full high-power (40x) field. To measure inhibitory postsynaptic potentials, current clamp recordings were performed while stimulating ChR2 using trains of light pulses (5 ms light pulses at 5 Hz, 20 Hz, and 40 Hz). In experiments in which glutamatergic and GABA_A_ receptors were blocked, drugs were bath applied at the following concentrations (in μM): 10 6-Cyano-7-nitroquinoxaline-2,3-dione disodium salt (CNQX, Tocris), 50 D-2-amino-5-phosphonopentanoic acid (D-APV) (Tocris), and 10 gabazine (Sigma). Drugs were prepared as concentrated stock solutions and were diluted in ACSF on the day of the experiment.

### Rule shift task

This cognitive flexibility task was described in^6^. Briefly, mice are singly-housed and habituated to a reverse light/dark cycle, and food intake is restricted until the mouse is 80-85% of the *ad libitum* feeding weight. At the start of each trial, the mouse was placed in its home cage to explore two bowls, each containing one odor and one digging medium, until it dug in one bowl, signifying a choice. As soon as a mouse began to dig in the incorrect bowl, the other bowl was removed, so there was no opportunity for “bowl switching.” (Digging is defined as the sustained displacement of the media within a bowl). The bait was a piece of a peanut butter chip (approximately 5-10 mg in weight) and the cues, either olfactory (odor) or somatosensory and visual (texture of the digging medium which hides the bait), were altered and counterbalanced. All cues were presented in small animal food bowls (All Living Things Nibble bowls, PetSmart) that were identical in color and size. Digging media were mixed with the odor (0.01% by volume) and peanut butter chip powder (0.1% by volume). All odors were ground dried spices (McCormick garlic and coriander), and unscented digging media (Mosser Lee White Sand Soil Cover, Natural Integrity Clumping Clay cat litter).

After mice reached their target weight, they underwent one day of habituation. On this day, mice were given ten consecutive trials with the baited food bowl to ascertain that they could reliably dig and that only one bowl contained food reward. All mice were able to dig for the reward. Mice do not undergo any other specific training before being tested on the task. Then, on Days 1 and 2 (and in some cases, on additional days as well), mice performed the task (this was the testing done for experiments). After the task was done for the day, the bowls were filled with different odor-medium combinations and food was evenly distributed among these bowls and given to the mouse so that the mouse would disregard any associations made earlier in the day.

Mice were tested through a series of trials. The determination of which odor and medium to pair and which side (left or right) contained the baited bowl was randomized (subject to the requirement that the same combination of pairing and side did not repeat on more than 3 consecutive trials) using http://random.org. On each trial, while the particular odor-medium combination present in each of the two bowls may have changed, the particular stimulus (e.g., a particular odor or medium) that signaled the presence of food reward remained constant over each portion of the task (initial association and rule shift). If the initial association paired a specific odor with food reward, then the digging medium would be considered the irrelevant dimension. The mouse is considered to have learned the initial association between stimulus and reward if it makes 8 correct choices during 10 consecutive trials. Each portion of the task ended when the mouse met this criterion. Following the initial association, the rule shift portion of the task began, and the particular stimulus associated with reward underwent an extra-dimensional shift. For example, if an odor had been associated with reward during the initial association, then a digging medium was associated with reward during the rule shift portion of the task. The mouse is considered to have learned this extra-dimensional rule shift if it makes 8 correct choices during 10 consecutive trials. When a mouse makes a correct choice on a trial, it is allowed to consume the food reward before the next trial. Following correct trials, the mouse is transferred from the home cage to a holding cage for about 10 seconds while the new bowls were set up (intertrial interval). After making an error on a trial, a mouse was transferred to the holding cage for about 2 minutes (intertrial interval). All animals performed the initial association in a similar number of trials (average: 10-15 trials). Experiments were performed blind to the virus injected. Videos were manually scored with a temporal resolution of 1 second.

For analyses (described below), the onset of digging was chosen as the time of a decision for two reasons. First, as noted above, once a mouse began to dig in the incorrect bowl, the other (correct) bowl was removed. Second, only upon the commencement of digging could a mouse determine whether reward was present in the chosen bowl and obtain feedback about whether or not it had made a correct choice.

### *In vivo* optogenetic stimulation

For behavioral experiments using optogenetic eNpHR stimulation: A 532 nm green laser (OEM Laser Systems, Inc.) was coupled to the mono fiber-optic cannula (Doric Lenses) with a zirconia sleeve (Doric Lenses) through a 200 μm diameter mono fiber-optic patch cord (Doric Lenses, Inc.) and adjusted such that the final light power was 2.5 mW.

Extended Data Fig. 1 shows experiments designed to control for potential behavioral effects of scattered light from one hemisphere activating eNpHR in PV neuron cell bodies in the contralateral hemisphere. These experiments used a final light power of 0.1 mW when connected to the 532 nm green laser (OEM Laser Systems, Inc.) or 638 nm red laser (Doric Lenses). To determine the appropriate light power for these experiments, a mouse was implanted with a dual fiberoptic cannula (Doric Lenses; DFC_200/240-0.22_2.3mm_GS0.7_FLT) without virus injection in order to measure light scattering from one mPFC to the contralateral hemisphere. Using a dual fiber-optic patch cord (Doric Lenses; DFP_200/240/900-0.22_2m_GS0.7-2FC), light was delivered to the mPFC on one side, and the light coming through the other side was measured using a light meter (Thor Labs, PM100D). The final light power delivered to one mPFC was 2.5 mW, across wavelengths 532 nm, 594 nm, and 638 nm – similar to what was used in the optogenetic and optogenetic + dual-site voltage indicator experiments. Measurements of 40 nW, 20 nW, and 50 nW at the contralateral fiber was observed, respectively. Accounting for transmission loss of the patch cord (e.g., only 80% of the light is transmitted from end to end), the scattered light power entering the fiber tip would be 50 nW, 25 nW, and 62.5 nW respectively. This measurement likely overestimates the actual light entering via the fiber tip located in brain parenchyma since it will include both scattered light within the brain and contamination from ambient room light. Conversely, this measurement only includes light located in the vicinity of the fiber tip that is traveling at an appropriate angle to enter the fiber (which has numerical aperture 0.22 implying a 25.4 degree acceptance angle). Therefore, to be extremely conservative, experiments in Extended Data Fig. 1 utilized a final light power of 0.1 mW, which is >1,000 times stronger than the measured scattered light power. Both 532 nm and 638 nm at this final light power ipsilateral to the viral injection site was used.

For all optogenetic stimulation behavior experiments, light stimulation began once mice reached the 80% criterion during the initial association portion of the task. Mice then performed three additional initial association trials with the light stimulation before the rule shift portion of the task began. The light stimulation did not alter the performance or behavior of the mice during these three extra trials of the initial association. Experiments were performed blind to virus injected.

### Combined dual-site voltage indicator imaging and optogenetic eNpHR stimulation

High-bandwidth, time-varying bulk fluorescence signals were measured at each recording site using the dual-site voltage-indicator technique described in^8^, with some modifications as described below.

### Optical apparatus

A fiber-optic stub (400 μm core, NA=0.48, low-autofluorescence fiber; Doric Lenses, MFC_400/430-0.48_2.8mm_ZF1.25_FLT) was stereotaxically implanted in each targeted brain region. A matching fiber-optic patch cord (Doric Lenses, MFP_400/430/1100-0.48_2m_FC-ZF1.25) provided a light path between the animal and a miniature, permanently-aligned optical bench, or ‘mini-cube’ (Doric Lenses, FMC5_E1(460-490)_F1(500-540)_E2(555-570)_F2(580-600)_S). One fiber on the ipsilateral side of the viral injection of either 1 uL of AAV2-EF1α-DIO-eNpHR3.0-BFP (Virovek) or 1 μL of AAV5-EF1α-DIO-mCherry (UNC Virus Core) was used to both deliver excitation light to and collect emitted fluorescence from that recording site. The fiber contralateral to the viral injection site was connected to a separate mini-cube (Doric Lenses, FMC6_E1(460-490)_F1(500-540)_E2(555-570)_F2(580-600)_O(628-642)_S) that was attached to a 638 nm laser (Doric Lenses) and was used to deliver excitation light and optogenetic stimulation to and collect emitted fluorescence from that recording site. The far end of the patch cord and each 1.25 mm diameter zirconia optical implant ferrule were cleaned with isopropanol before each recording, then securely attached via a zirconia sleeve.

For the first mini-cube sans laser port, optics allow for the simultaneous monitoring of two spectrally-separated fluorophores, with dichroic mirrors and cleanup filters chosen to match the excitation and emission spectra of the voltage sensor and reference fluorophores in use (‘mNeon’ voltage sensor channel: Ex. 460-490 nm, Em. 500-540 nm; ‘Red’ control fluorophore: Ex. 555-570 nm, Em. 580-600 nm). The mini-cube optics are sealed and permanently aligned and all 5 ports (sample to animal, 2 excitation lines, 2 emission lines) are provided with matched coupling optics and FC connectors to allow for a modular system design.

For the second mini-cube that contains a port for optogenetic stimulation, a 565 nm LED and 555-570 nm filter is used to excite tdTomato, a 580-600 nm filter is used for tdTomato emission, and a 638 nm laser with 628-642 nm filter is used to excite eNpHR. The light used to excite eNpHR does not interfere with the genetically-encoded voltage indicator based measurements of synchrony, and the light used to excite tdTomato does not activate eNpHR enough to affect rule shift performance. This is due to two factors. First, the eNpHR and tdTomato excitation spectra are offset, e.g., at 605 nm, eNpHR activation is near its peak whereas relative excitation of tdTomato is <1%. Second, the intensity used to excite tdTomato is just ∼0.1 mW, which is ∼50-fold less than what was used to activate eNpHR and disrupt rule shift performance. The mini-cube optics are sealed and permanently aligned and all 6 ports (sample to animal, 3 excitation lines, 2 emission lines) are provided with matched coupling optics and FC connectors to allow for a modular system design.

To perform dual-site voltage indicator recordings, excitation light for each of the two color channels was provided by a fiber-coupled LED (Center wavelengths 490 nm and 565 nm, Thorlabs M490F3 and M565F3) connected to the mini-cube by a patch cord (200 μm, NA=0.39; Thorlabs M75L01). Using a smaller diameter for this patch cord than for the patch cord from the cube to the animal is critical to reduce the excitation spot size on the output fiber face and thus avoid cladding autofluorescence. LEDs were controlled by a 4-channel, 10kHz-bandwidth current source (Thorlabs DC4104). LED current was adjusted to give a final light power at the animal (averaged during modulation, see below) of approximately 200 μW for the mNeon channel (460-490 nm excitation), and 100 μW for the Red channel (555-570 nm excitation).

Each of the two emission ports on the mini-cube was connected to an adjustable-gain photoreceiver (Femto, Berlin, Germany, OE-200-Si-FC; Bandwidth set to 7kHz, AC-coupled, ‘Low’gain of ∼5×10^7 V/W) using a large-core high-NA fiber to maximize throughput (600 μm core, NA=0.48 (Doric lenses, MFP_600/630/LWMJ-0.48_0.5m_FC-FC).

Note that, for dual-site voltage indicator recordings and optogenetics experiments, two completely independent optical setups were employed, with separate implants, patch cords, mini-cubes, LEDs, a separate laser, photoreceivers, and lock-in amplifiers.

### Modulation and lock-in detection

At each recording site, each of the two LEDs was sinusoidally modulated at a distinct carrier frequency to reduce crosstalk due to overlap in fluorophore spectra. The corresponding photoreceiver outputs were then demodulated using lock-in amplification techniques. A single instrument (Stanford Research Systems, SR860) was used to generate the modulation waveform for each LED and to demodulate the photoreceiver output at the carrier frequency. To further reduce cross-talk between recording sites, distinct carrier frequencies (2, 2.5, 3.5 and 4 kHz) were used across sites. Low-pass filters on the lock-in amplifiers were selected to reject noise above the frequencies under study (cascade of 4 Gaussian FIR filters with 84 Hz equivalent noise bandwidth; final attenuation of signals are approximately −1dB (89% of original magnitude) at 20 Hz, −3dB (71% of original magnitude) at 40 Hz, and −6dB (50% of original magnitude) at 60 Hz).

### *In vivo* dual-site voltage indicator imaging

Analog signals were digitized by a multichannel real-time signal processor (Tucker-Davis Technologies, Alachua, Florida; RX-8). The commercial software Synapse (Tucker-Davis Technologies) running on a PC was used to control the signal processor, write data streams to disk, and to record synchronized video from a generic infrared USB webcam (Ailipu Technology, Shenzhen, China, ELP-USB100W05MT-DL36). Lock-in amplifier outputs were digitized at 3 kHz.

### Combined dual-site voltage indicator imaging and optogenetics analysis

Analysis of voltage indicator data was described previously in (*17*) and was facilitated using the signal processing toolbox and MATLAB (Mathworks), using the following functions: fir1, filtfilt, and regress. All four signals during the entire time series of the experiment (left mNeon, left tdTomato, right mNeon, right tdTomato) were first filtered around a frequency of interest. To quantify zero-phase lag cross-hemispheric synchronization between left and right mNeon signals, a linear regression analysis was performed to predict the right mNeon signal using the following inputs: left mNeon signal, left tdTomato signal, and right tdTomato signal. The goodness of fit is compared to how well the regression works if the left mNeon signal is shuffled, i.e., if a randomly chosen segment of the original left mNeon signal is used, instead of the segment recorded at the same time as the right mNeon signal. *R*^*2*^ values are calculated as a function of time using one second segments and compared to the 99^th^ percentile of the distribution of *R*^*2*^ values obtained from 100 fits to randomly-shuffled data. The fraction of timepoints at which the *R*^*2*^ obtained from actual data exceeds the 99^th^ percentile of the *R*^*2*^ values obtained from shuffled data was used to measure zero-phase lag synchronization between the left and right mNeon signals.

This analysis was performed at the time of the decision (e.g., immediately following the beginning of digging in one bowl, until the end of digging), and smoothed measurements over a 5 minute time window following the time point of interest. The first five trials of the rule shift were analyzed. Experiments were performed, scored, and analyzed blind to virus injected.

### Combined calcium imaging and optogenetics

Imaging data were collected using a miniaturized one-photon microscope (nVoke2; Inscopix Inc.). GCaMP7f signals (calcium activity) were detected using 435–460 nm excitation LED (0.1–0.2 mW), and optogenetic stimulation of eNpHR expressing axons was performed using a second 590-650 nm excitation LED (1–2 mW light power). nVoke2 software (Inscopix) was used to control the microscope and collect imaging data. Images were acquired at 20 frames per second, spatially downsampled (4x), and were stored for offline data processing. An input TTL from a separate ANY-maze computer (Stoelting Europe) to the nVoke2 acquisition software were used to synchronize calcium imaging and mouse behavior movies.

### Combined calcium imaging and optogenetics analysis

Calcium imaging movies were preprocessed using Inscopix Data Processing Software (IDPS; Inscopix, Inc.). The video frames were spatially filtered (band-pass) with cut-offs set to 0.008 pixel^-1^ (low) and 0.3 pixel^-1^ (high) followed by frame-by-frame motion correction for removing movement artifacts associated with respiration and head-unrestrained behavior. The mean image over the imaging session was computed, and the *dF/F* was computed using this mean image. The resultant preprocessed movies were then exported into MATLAB, and cell segmentation was performed using an open-source calcium imaging software (CIAPKG)^29^. Specifically, a Principal Component Analysis/Independent Component Analysis (PCA/ICA) approach was used to detect and extract ROIs (presumed neurons) per field of view^30^. For each movie, the extracted output neurons were then manually sorted to remove overlapping neurons, neurons with low SNR, and neurons with aberrant shapes. Accepted neurons and their calcium activity traces were exported to MATLAB for further analysis using custom scripts as previously described in^31^. Briefly, the standard deviation (σ) of the calcium movie was calculated and this was used to perform threshold-based event detection on the traces by first detecting increases in *dF/F* exceeding 2 (over one second). Subsequently, events were detected that exceeded 10 for over two seconds and had a total area under the curve higher than 150σ. The peak of the event was estimated as the local maximum of the entire event. For an extracted output neuron, active frames were marked as the period from the beginning of an event until the calcium signal decreased 30% from the peak of the event (up to a maximum of 2 seconds).

We calculated the similarity of population activity vectors using the ‘cosine similarity,’ which is equivalent to computing the normalized dot product between the two vectors, i.e.:

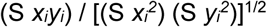

where *x*_*i*_ and *y*_*i*_ represent the average activity of the *i*th neuron in the two population activity vectors. Each vector was the average of activity during the 10 seconds immediately following a choice. We computed the similarity between each pair of vectors, then averaged this similarity across all the pairs from one mouse.

The first five trials of the rule shift were analyzed. For experiments in Fig. 1G–H, some mice did not make an incorrect decision in the first five initial association trials; therefore, the first error trial that occurred during the initial association was used for analysis. For Extended Data Fig. 5C, the last five initial association trials and the additional initial association trials during optogenetic inhibition were analyzed.

### Histology and imaging

All mice used for behavioral and imaging experiments were anesthetized with Euthasol and transcardially perfused with 30 mL of ice-cold 0.01 M PBS followed by 30 ml of ice-cold 4% paraformaldehyde (PFA) in PBS. Brains were extracted and stored in 4% PFA for 24 hours at 4 °C before being stored in PBS. 50 μm and 70 μm slices were obtained on a Leica VT100S and mounted on slides. All imaging was performed on an Olympus MVX10, Nikon Eclipse 90i, Zeiss LSM510, and Zeiss Axioskop2. All mice were verified to have virus-driven expression and optical fibers located in the mPFC.

### Data analyses and statistics

Statistical analyses were performed using Prism 8 (GraphPad) and detailed in the corresponding figure legends. Quantitative data are expressed as the mean and error bars represent the standard error of the mean (SEM). Group comparisons were made using two-way ANOVA followed by Bonferroni post-hoc tests to control for multiple comparisons. Paired and unpaired two-tailed Student’s *t*-tests were used to make single-variable comparisons. Similarity of variance between groups was confirmed by the *F* test. Measurements were taken from distinct samples and from samples that were measured repeatedly. *P* = *, < 0.05; **, < 0.01; ***, < 0.001; ****, < 0.0001. Comparisons with no asterisk had *P* > 0.05 and were considered not significant. No statistical methods were used to pre-determine sample sizes but our sample size choice was based on previous studies (*13, 17*) and are consistent with those generally employed in the field. Data distribution was assumed to be normal, but this was not formally tested.

## Acknowledgments

V.S.S. is supported by grants from the US National Institute of Health (R01MH121342 and R01NS116594), McKnight Memory and Cognitive Disorders, and the Brain Research Foundation. K.K.A.C. is supported by the Institut National de la Santé et de la Recherche Médicale (Inserm) and Marie Skłodowska-Curie Individual Fellowship (MSCA-IF).

## Author contributions

K.K.A.C. and V.S.S. designed the study. K.K.A.C. conducted all experiments and analyses, with the exception of Extended Data Fig. 1E–1H, which was performed by J.S. K.K.A.C. and V.S.S. wrote the paper.

## Supplementary information

is available for this paper.

## Competing interests

The authors declare no competing interests.

## Data and materials availability

The data that support the findings of this study are available from the corresponding author upon reasonable request, and will be deposited in a public repository prior to publication.

## Code availability

Custom codes for analysis and modelling were written in MATLAB and are available from the corresponding author upon request.

**Extended Data Fig. 1:**
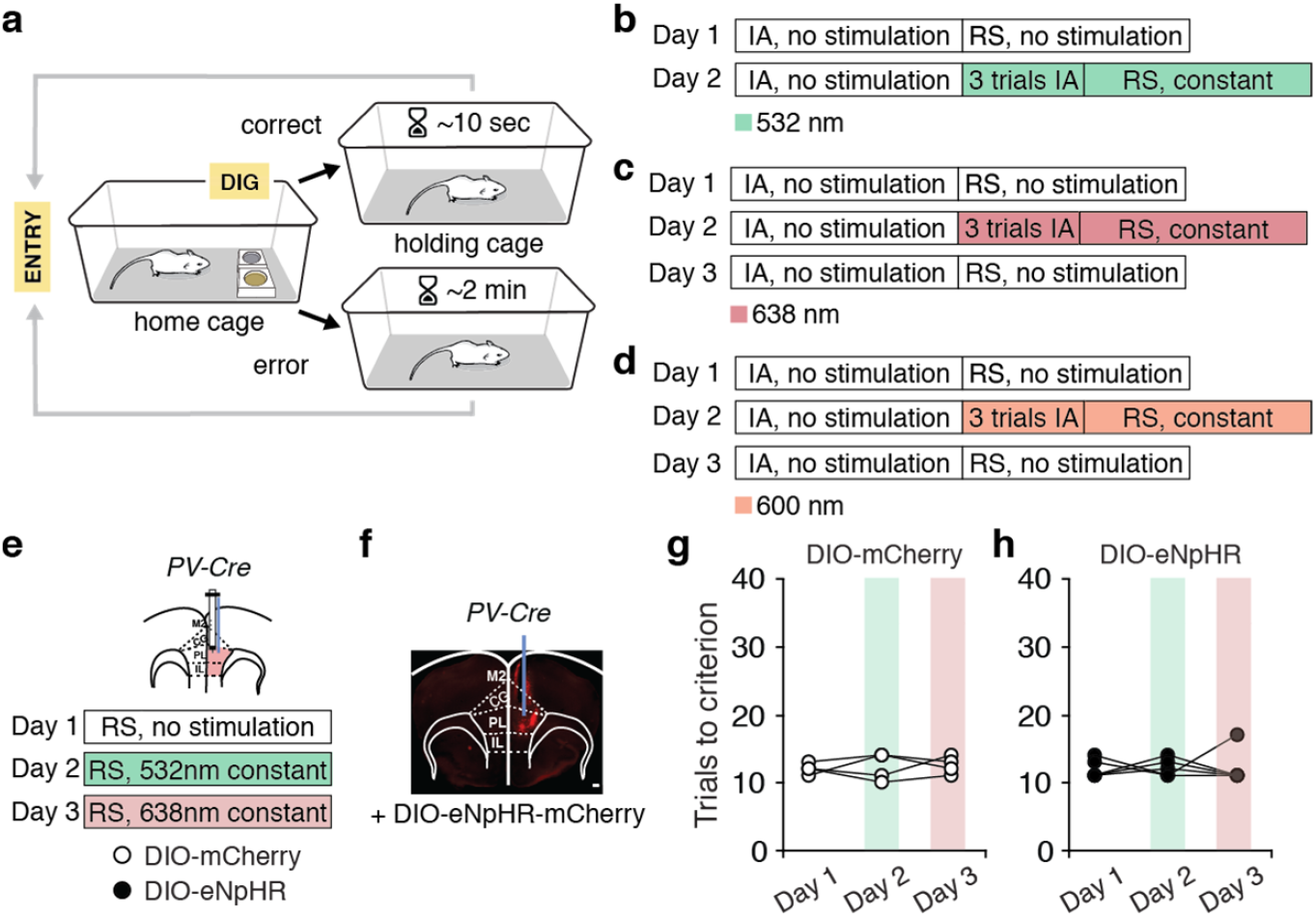
Rule shift task description and control experiments using weak light delivery. **a**, Trial timeline. A mouse begins each trial when it is placed in the home cage, then makes a decision, indicated by digging in one bowl. If the mouse is correct, the food reward is consumed. The mouse is then transferred to the holding cage until the next trial. The intertrial interval (ITI) is much longer after errors. **b**, For optogenetic inhibition behavior experiments in **Fig. 2**, Day 1: no light delivery during the initial association (IA), nor during the rule shift (RS); Day 2: no light is delivered while mice learn the IA, but once mice meet the criterion for learning (8/10 consecutive trials correct), we begin delivering continuous 532 nm light and test the animal for 3 additional IA trials, before switching to the RS portion of the task. **c**, For optogenetic inhibition dual-site voltage indicator experiments in **Fig. 3**, Day 1: no light delivery during the IA, nor during the RS; Day 2: no light delivery during the IA, but continuous light delivery of 638 nm begins during 3 additional IA trials, followed by the RS; Day 3: no light delivery during the IA, nor during the RS. **d**, For optogenetic inhibition + microendoscopic calcium imaging experiments in **Fig. 4**, Day 1: no light delivery during the IA, nor during the RS; Day 2: no light delivery during the IA, but continuous light delivery of 600 nm begins during 3 additional IA trials, followed by the RS; Day 3: no light delivery during the IA, nor during the RS. **e**, Control experiments to verify that weak light delivery does not affect rule shift learning. Experimental design: Day 1, no light delivery; Day 2, continuous 0.1 mW 532 nm light is delivered during the RS; Day 3, continuous light at 0.1 mW 638 nm for inhibition during the RS. **f**, Representative image showing AAV-DIO-eNpHR-mCherry (DIO-eNpHR) expression and a fiber-optic cannula in the same mPFC hemisphere in a *PV-Cre* mouse. Scale bar, 250 μm. **g, h**, Performance in the rule shift task did not vary across days in either controls, or DIO-eNpHR-injected mice (two-way ANOVA (task day × virus); interaction: *F*_(2,14)_ = 0.01721, *P* = 0.983, *n* = 4 control mice, 5 eNpHR mice). Two-way ANOVA followed by Bonferroni post hoc comparisons was used.

**Extended Data Fig. 2:**
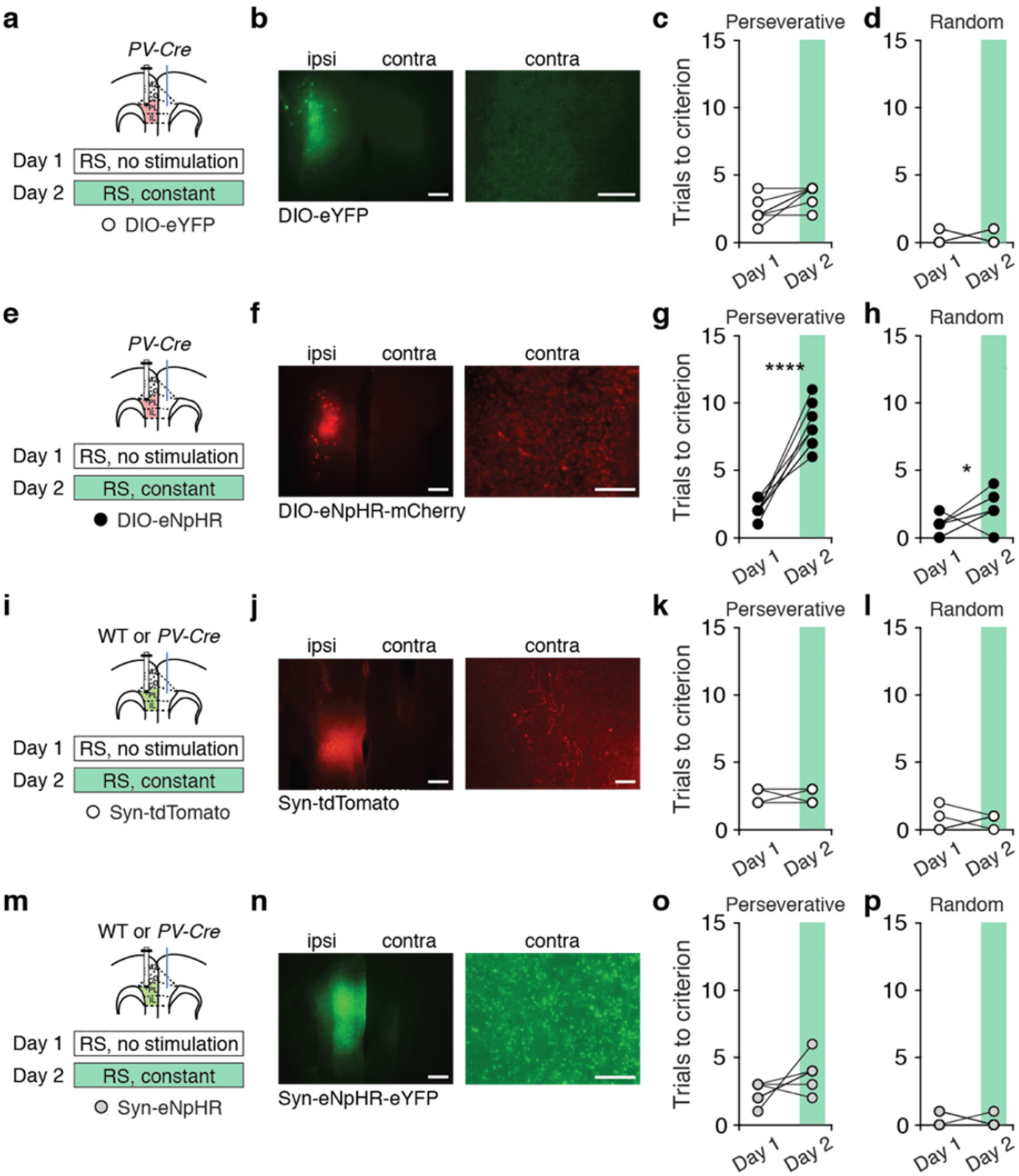
Optogenetic inhibition of callosal PV terminals (but not nonspecific inhibition of all callosal terminals) increases errors during rule shifts. **a, e, i, m**, Experimental design: Day 1, no light delivery; Day 2, continuous light for optogenetic inhibition of callosal PV terminals during the rule shift (RS). **b, f, j, n**, Representative image showing viral expression in the mPFC ipsilateral to the injection (ipsi), and labeled callosal terminals in the contralateral mPFC (contra). Scale bars, 250 μm and 100 μm, respectively. **c, g**, Optogenetic inhibition of callosal PV terminals increases perseverative errors in DIO-eNpHR-expressing mice (*n* = 8 mice) compared to DIO-eYFP-expressing controls (*n* = 7 mice; two-way ANOVA; main effect of task day: *F*_(1,13)_ = 73.42, *P* < 0.0001; main effect of virus: *F*_(1,13)_ = 37.06, *P* < 0.0001; interaction: *F*_(1,13)_ = 31.31, *P* < 0.0001). **d, h**, Optogenetic inhibition of callosal PV terminals has a marginal effect on random errors (two-way ANOVA (task day × virus); interaction: *F*_(1,13)_ = 4.617, *P* = 0.0511). **c, d**, Light delivery does not affect the number of perseverative (post hoc *t*_(13)_ = 2.036, *P* = 0.1254) or random (post hoc *t*_(13)_ = 0.0, *P* > 0.9999) errors in DIO-eYFP-expressing controls. **g, h**, Optogenetic inhibition of callosal PV terminals on Day 2 increased the number of perseverative (post hoc *t*_(13)_ = 10.37, *P* < 0.0001) and random (post hoc *t*_(13)_ = 3.145, *P* = 0.0155) errors compared to no stimulation on Day 1. **k, l, o, p**, Optogenetic inhibition of all callosal projections has no effect on perseverative errors in Syn-eNpHR-expressing mice (*n* = 6) compared to controls (*n* = 4; two-way ANOVA (task day × virus); interaction: *F*_(1,8)_ = 0.0, *P* > 0.9999) nor on random errors (two-way ANOVA (task day × virus); interaction: *F*_(1,8)_ = 0.07805, *P* = 0.787). Two-way ANOVA followed by Bonferroni post hoc comparisons was used. *\*P* < 0.05, \*\*\*\**P* < 0.0001.

**Extended Data Fig. 3:**
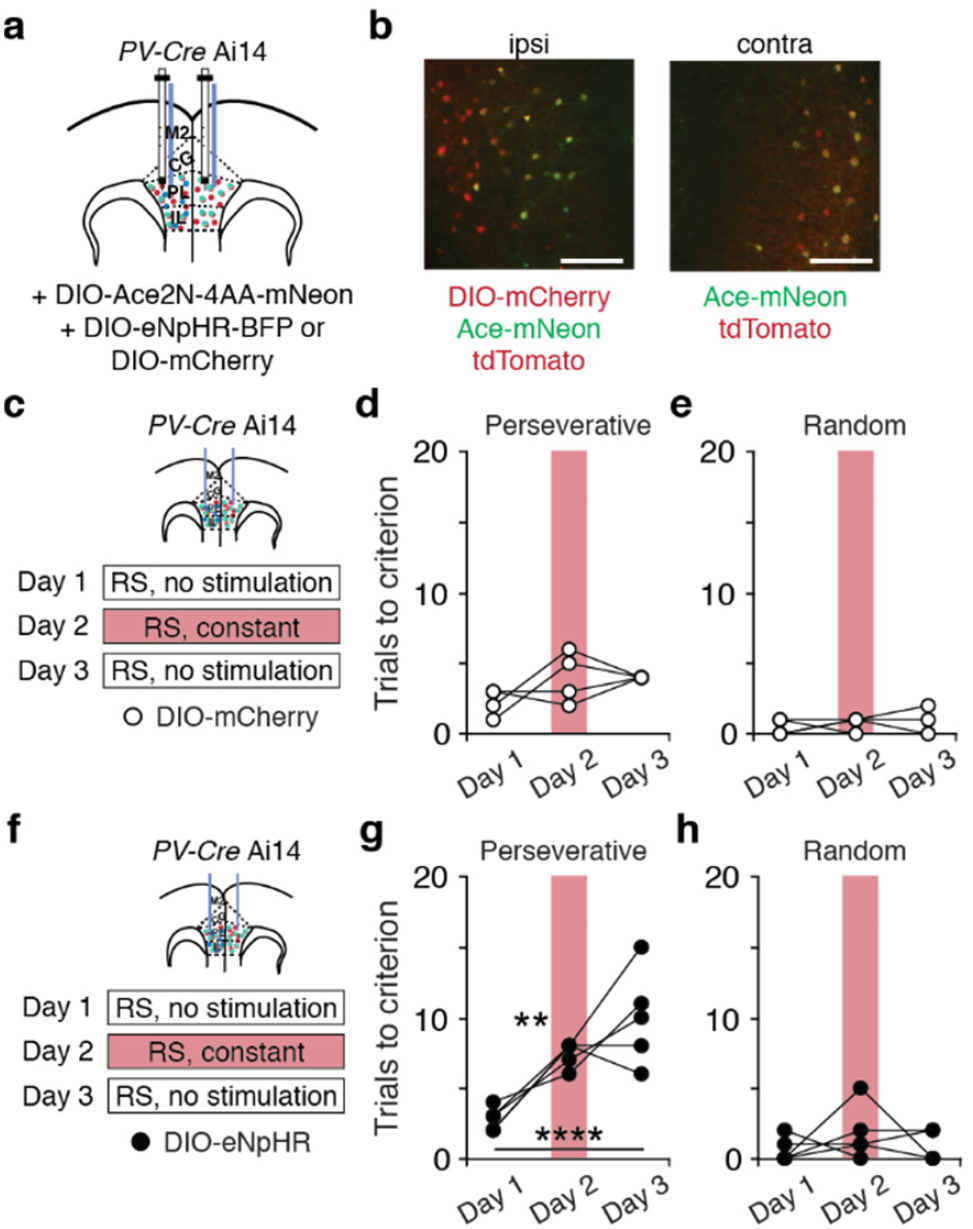
Optogenetic inhibition of callosal PV terminals, delivered while measuring signals from voltage indicators, increases errors during rule shifts (RS). **a**, *PV-Cre* Ai14 mice had bilateral AAV-DIO-Ace2N-4AA-mNeon (Ace-mNeon) injections, an ipsilateral AAV-DIO-eNpHR-BFP (DIO-eNpHR) or AAV-DIO-mCherry injection and multimode fiber-optic implants in both prefrontal cortices. **b**, Representative images from mice injected with a control virus (DIO-mCherry), showing mCherry, Ace-mNeon, and tdTomato expression in the mPFC ipsilateral to the virus injection (ipsi), and Ace-mNeon and tdTomato in the contralateral hemisphere (contra). Scale bars, 50 μm. **c, f**, Experimental design: Day 1, no light delivery; Day 2, continuous light for inhibition during the rule shift (RS); Day 3, no light delivery. **d, e, g, h**, Optogenetic inhibition of callosal PV terminals increases perseverative errors in DIO-eNpHR-expressing mice (*n* = 5 mice, **g-h**) compared to DIO-mCherry-expressing controls (*n* = 4 mice, **d-e**; two-way ANOVA (task day × virus); interaction: *F*_(2,14)_ = 5.226, *P* = 0.0202), but has no effect on random errors (two-way ANOVA (task day × virus); interaction: *F*_(2,14)_ = 0.4552, *P* = 0.6434). **d, e**, Light delivery does not affect the number of perseverative (Day 1 to Day 2: post hoc *t*_(14)_ = 1.392, *P* = 0.5566; Day 1 to Day 3: post hoc *t*_(14)_ = 1.392, *P* = 0.5566; Day 2 to Day 3: post hoc *t*_(14)_ = 0.0, *P* > 0.9999) or random (Day 1 to Day 2: post hoc *t*_(14)_ = 0.284, *P* > 0.9999; Day 1 to Day 3: post hoc *t*_(14)_ = 0.284, *P* > 0.9999; Day 2 to Day 3: post hoc *t*_(14)_ = 0.0, *P* > 0.9999) errors in controls across days. **g, h**, Optogenetic inhibition of callosal PV terminals induces perseveration on Day 2 and Day 3 compared to no stimulation on Day 1 (Day 1 to Day 2: post hoc *t*_(14)_ = 4.092, *P* = 0.0033; Day 1 to Day 3: post hoc *t*_(14)_ = 6.405, *P* < 0.0001; Day 2 to Day 3: post hoc *t*_(14)_ = 2.313, *P* = 0.1094), but has no effect on random errors (Day 1 to Day 2: post hoc *t*_(14)_ = 1.524, *P* = 0.4493; Day 1 to Day 3: post hoc *t*_(14)_ = 0.254, *P* > 0.9999; Day 2 to Day 3: post hoc *t*_(14)_ = 1.27, *P* = 0.6744). Two-way ANOVA followed by Bonferroni post hoc comparisons was used. *\*\*P* < 0.01, \*\*\*\**P* < 0.0001.

**Extended Data Fig. 4:**
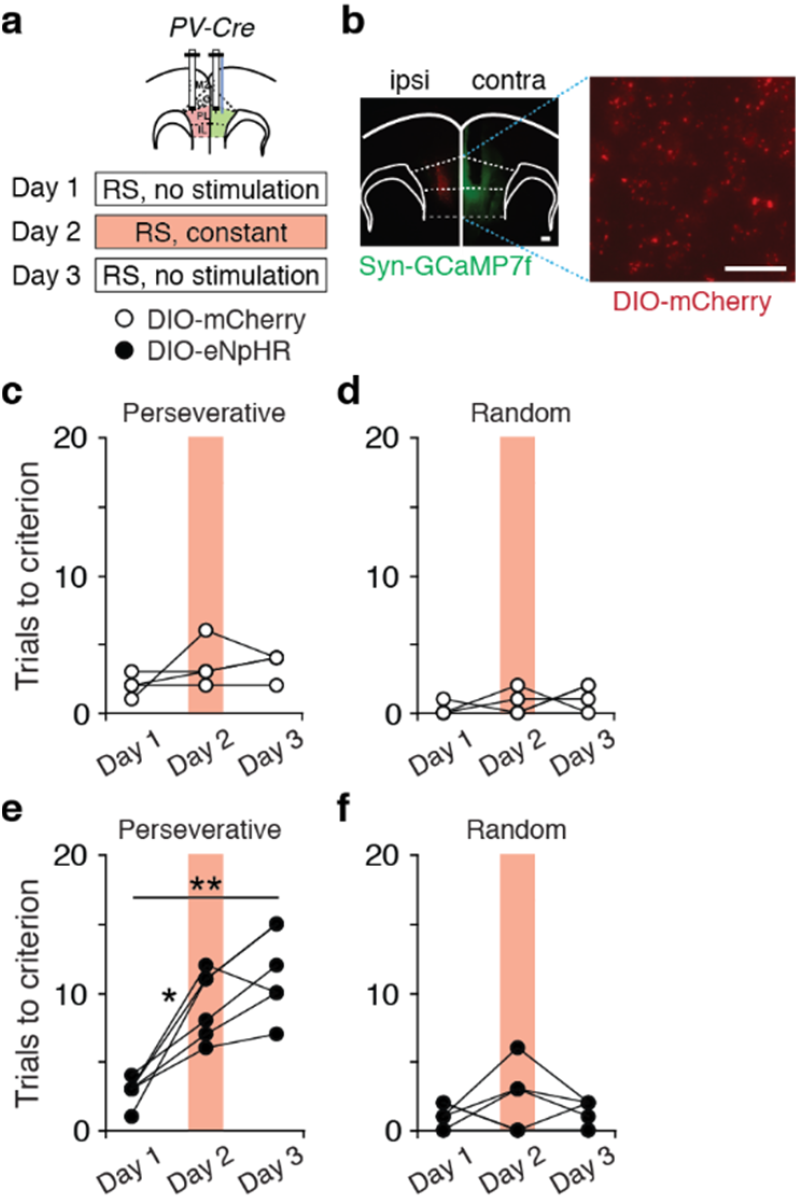
Optogenetic inhibition of callosal terminals, delivered during concomitant calcium imaging, increases errors during rule shifts (RS). **a**, *PV-Cre* mice had AAV-DIO-eNpHR-mCherry (DIO-eNpHR) or a control virus (AAV-DIO-mCherry) injected in one mPFC and AAV-Synapsin-GCaMP7f (Syn-GCaMP7f) injected in the contralateral hemisphere. A GRIN lens, connected to a miniscope, was implanted in contralateral to the site of eNpHR / control virus injection. Experimental design: Day 1, no light delivery for optogenetic inhibition; Day 2, continuous light for inhibition during the rule shift (RS); Day 3, no light delivery for optogenetic inhibition (light was delivered for calcium imaging on all days). **b**, Left: Representative image showing AAV-DIO-mCherry (DIO-mCherry) injected in the ipsilateral mPFC (ipsi) and Syn-GCaMP7f in the contralateral hemisphere (contra). Right: DIO-mCherry expression in callosal PV axonal fibers. Scale bars, 100 μm and 50 μm, respectively. **c, e**, Optogenetic inhibition of callosal PV terminals increases perseverative errors in DIO-eNpHR-expressing mice (*n* = 6 mice, **c**) compared to DIO-mCherry-expressing controls (*n* = 4 mice, **d**; two-way ANOVA(task day × virus); interaction: *F*_(2,16)_ = 9.054, *P* = 0.0023), but has no effect on random errors (two-way ANOVA (task day × virus); interaction: *F*_(2,16)_ = 0.5463, *P* = 0.5895). **c, d**, Light delivery does not affect the number of perseverative (Day 1 to Day 2: post hoc *t*_(3)_ = 1.26, *P* = 0.8901; Day 1 to Day 3: post hoc *t*_(3)_ = 2.324, *P* = 0.3082; Day 2 to Day 3: post hoc *t*_(3)_ = 0.0, *P* > 0.9999) or random (Day 1 to Day 2: post hoc *t*_(3)_ = 0.7746, *P* > 0.9999; Day 1 to Day 3: post hoc *t*_(3)_ = 2.449, *P* > 0.2752; Day 2 to Day 3: post hoc *t*_(3)_ = 0.5222, *P* > 0.9999) errors in DIO-mCherry-expressing controls across days. **e, f**, Optogenetic inhibition of callosal PV terminals increases the number of perseverative errors on Day 2 and Day 3 compared to no stimulation on Day 1 (Day 1 to Day 2: post hoc *t*_(5)_ = 5.152, *P* = 0.0108; Day 1 to Day 3: post hoc *t*_(5)_ = 5.785, *P* = 0.0065; Day 2 to Day 3: post hoc *t*_(5)_ = 2.36, *P* = 0.1943), but has no effect on random errors (Day 1 to Day 2: post hoc *t*_(5)_ = 0.866, *P* > 0.9999; Day 1 to Day 3: post hoc *t*_(5)_ = 0.2225, *P* > 0.9999; Day 2 to Day 3: post hoc *t*_(5)_ = 1.0, *P* > 0.9999). Two-way ANOVA followed by Bonferroni post hoc comparisons was used. *\*P* < 0.05, \*\**P* < 0.01.

**Extended Data Fig. 5:**
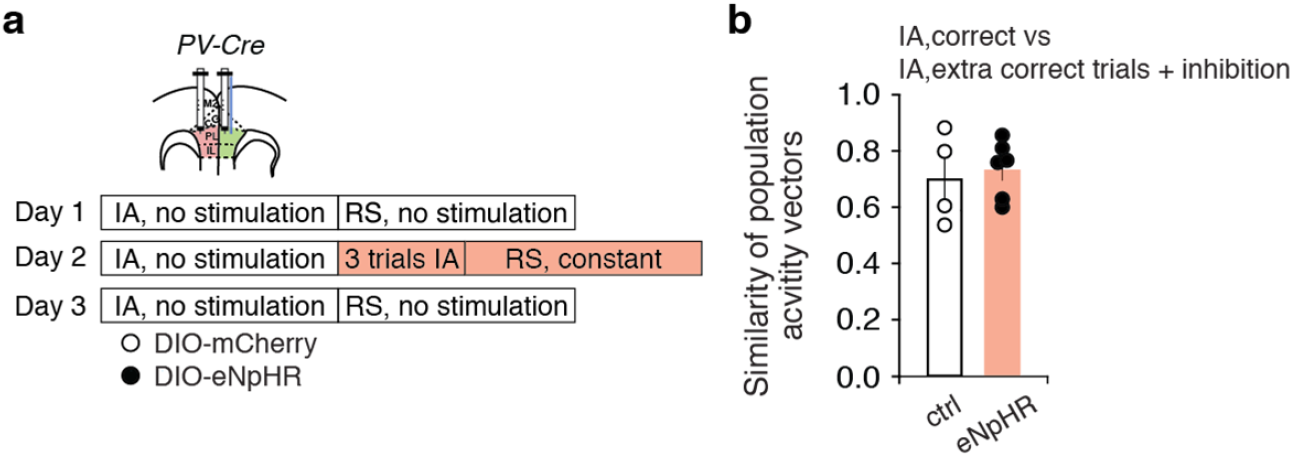
Optogenetic inhibition of callosal PV terminals does not affect the similarity of population activity vectors across groups on correct trials before the rule shift. **a**, *PV-Cre* mice had AAV-DIO-eNpHR-mCherry (DIO-eNpHR) or a control (ctrl) virus (AAV-DIO-mCherry) injected into one mPFC, and AAV-Synapsin-GCaMP7f injected and a GRIN lens connected to a miniscope implanted in the contralateral hemisphere. Experimental design: Day 1, no light delivery for optogenetic inhibition; Day 2, continuous light for optogenetic inhibition during the rule shift (RS); Day 3, no light delivery for optogenetic inhibition (light was delivered for calcium imaging on all days). **b**, On Day 2, after mice reach the learning criterion for the IA, we began delivering optogenetic inhibition and performed three additional IA trials, followed by the rule shift. Optogenetic inhibition of callosal PV terminals did not affect the similarity between correct trial activity vectors during these three additional IA trials and the preceding five IA trials (two-tailed, unpaired *t*-test, *t*_(8)_ = 0.3812, *P* = 0.7129). Thus, the effect of inhibiting callosal PV terminals to suppress changes in activity does not occur prior to the rule shift.

## Notes

### Competing Interest Statement

The authors have declared no competing interest.

